# Genetic dissection of the obesity paradox in carotid atherosclerosis using a hyperlipidemic mouse cohort

**DOI:** 10.64898/2026.08.13.744610

**Authors:** Kiyan Parvaresh, Firas Dalloul, Mei-Hua Chen, Lisa J. Shi, Muhammad Sarfraz Ali, Hideyuki Torikai, Weibin Shi

**Affiliations:** Department of Radiology and Medical Imaging, Charlottesville, VA 22908; Biochemistry & Molecular Genetics, University of Virginia, Charlottesville, VA 22908

## Abstract

**Background:** Overweight and obese individuals often exhibit lower mortality rates or better prognoses than lean or normal-weight individuals with stroke and other diseases, a phenomenon called the “obesity paradox”. Carotid atherosclerosis is the primary cause of ischemic stroke, and body weight serves as a reliable surrogate for adiposity in mice.

**Methods:** Phenotypic and genetic connections of carotid atherosclerosis with body weight were evaluated in 299 F2 mice derived from BALB/cJ and LP/J *Apoe* knockout (*Apoe^-/-^*) mice. F2 mice were fed a Western diet for 12 weeks. Atherosclerotic lesion sizes in left carotid arteries, body weight, coat color, plasma lipids, glucose, small dense LDL ApoB, and malondialdehyde were measured, and 11,000 single nucleotide polymorphism (SNP) markers were genotyped.

**Results:** Carotid lesion sizes inversely correlated with body weight in both sexes. Genome-wide scans identified two significant quantitative trait loci (QTLs) for carotid atherosclerosis on chromosomes (Chr) 6 and 15 in an additive sex model, and five QTLs on Chr 6, 7, 12, 13, and 15 in an interactive sex model. Adjusting for body weight variation downgraded Chr 15 QTL (*Cath5*) in both models, whereas other QTLs upgraded in the additive sex model and downgraded in the interactive sex model. Human syntenic region of *Cath5* associated with carotid intima-medial thickness (cIMT) and waist-to-hip ratio (WHR).

**Conclusions:** These findings indicate that the obesity paradox in carotid atherosclerosis is partially driven by shared genetic components that exert opposing effects on adiposity and plaque development and act through sex-dependent mechanisms.

## Introduction

Stroke is the leading cause of adult disability and the fourth leading cause of death in the United States ^1^. Ischemic stroke, which occurs when blood flow to the brain is obstructed, accounts for over 80% of all stroke cases ^2^. A large portion of ischemic strokes stem from atherosclerosis in the carotid arteries, where plaque buildup and rupture narrow the arterial lumen and block cerebral blood flow ^3^. Atherosclerosis is a complex trait influenced by environmental factors and strong genetic components, with heritability estimated at ∼50% for carotid plaque and carotid intima-media thickness (cIMT) ^4,5,6^, two validated ultrasound surrogates of subclinical atherosclerosis. A recent meta-analysis of seven genome-wide association studies (GWAS) involving 131,000 individuals across diverse ancestries identified 59 independent loci for cIMT ^7^. However, because the individual effect sizes of human GWAS-detected variants are typically small, pinpointing causal genes remains challenging. Consequently, parallel quantitative genetic approaches using animal models are essential to accelerate gene discovery in carotid atherosclerosis.

*Apoe* knockout (*Apoe*^-/-^) mice develop all phases of atherosclerotic lesions in large and medium-sized arteries, which are heavily influenced by diet and genetic backgrounds ^8^. Using segregating F2 populations from *Apoe*^-/-^ mouse strains across various genetic backgrounds, we and others conduct quantitative trait locus (QTL) analysis for carotid atherosclerosis. To date, 20 significant QTLs for carotid atherosclerosis have been identified from five independent crosses of *Apoe*^-/-^ strains ^9,10,11,12,13^. Crucially, segregating F2 cohorts also enables the simultaneous dissection of phenotypic correlations and shared genetic architecture between distinct traits ^14,15,16^.

Obesity, characterized by excessive adiposity, is an established risk factor for stroke. However, clinical observations reveal that established stroke patients with higher body mass index (BMI) frequently correlates with lower mortality rates and improved long-term prognosis ^17,18,19,20,21^. This phenomenon, termed the “obesity paradox”, often manifests as a U-shaped or inverse relationship, where normal-weight or mildly overweight individuals experience better outcomes than underweight or severely obese counterparts ^22,23^.

Whether the obesity paradox reflects genuine biological protection, such as elevated protective adipokines or greater metabolic reserves, or merely methodological artifacts, like the inability of BMI to distinguish fat from lean muscle mass, remains intensely debated. Disentangling these mechanisms in human populations is difficult due to environmental confounders, dietary variations, and complex genetic heterogeneity. In contrast, mouse models allow for strict diet and environmental control and genetic manipulation. Total body weight serves as a validated, highly reliable surrogate for adiposity, enabling precise quantitative genetic mapping ^16^. Moreover, hyperlipidemic *Apoe^-/-^*mice recapitulate key elements of both metabolic dysfunction and plaque development, offering an ideal system to interrogate how body mass and atherosclerosis interact at the genome level. Here, we evaluated the phenotypic and genetic connections between body weight and carotid atherosclerosis in a segregating F2 cohort derived from BALB/cJ and LP/J *Apoe^-/-^*mice. Through high-density SNP genotyping and quantitative trait mapping, we sought to determine whether shared genetic loci with opposing effects on body weight and plaque burden contribute to the biological mechanisms underlying the obesity paradox.

## Methods

### Mice

An F2 cohort was generated from an intercross between male LP/J (LP) and female BALB/cJ (BALB) *Apoe*^-/-^ mice, as previously reported ^16^. At 6 weeks of age, all F2 mice started a 12-week Western diet (TD 88137, Envigo). Non-fasting blood samples were collected twice: once before the diet began and once after 11 weeks on the diet. At the end of the 12-week feeding period, body weight was measured and fasting blood samples were collected following an overnight fast. All blood samples were taken from the retro-orbital plexus of mice under isoflurane anesthesia, and plasma was prepared as described previously ^24^. All procedures were approved by the Institutional Animal Care and Use Committee (protocol #: 3109).

### Phenotypic analyses

Atherosclerotic lesion sizes in the left carotid artery of F2 mice were measured as reported ^9^. Briefly, the vasculature was perfusion-fixed with 10% formalin via the left ventricle. The distal portion of the common carotid artery and adjacent branches were dissected and embedded in Tissue-Tek optimum cutting (OCT) compound. 10-μm-thick cryosections were collected every three sections and stained with oil red O and hematoxylin. Lesion sizes were quantified using Zen imaging software. For each mouse, the mean lesion area was calculated from the five sections exhibiting the largest lesions and used for statistical analysis.

Plasma concentrations of cholesterol and triglycerides were measured using enzymatic assays as reported ^24^. HDL cholesterol was measured after precipitation of other lipoproteins using a phosphotungstate-magnesium reagent (Wako Diagnostics). Non-HDL cholesterol was calculated by subtracting HDL cholesterol from total cholesterol concentrations. Plasma glucose was measured using a Sigma assay kit (Cat. # GAHK20) as reported ^24^. Plasma small dense LDL was determined by first precipitating non-small dense LDL, IDL, VLDL lipoproteins, and chylomicrons using a phosphotungstate-magnesium reagent (FUJIFILM Wako) and then measuring the remaining ApoB concentration using an ELISA kit (MyBiosource, San Diego, CA, USA; Cat. #: MBS937790) ^25^. Plasma levels of malondialdehyde, an indicator of lipid peroxidation, were measured using a Cayman Thiobarbituric Acid Reactive Substances assay kit (TBARS) kit (Cat. # 10009055).

### Genotypic analysis

DNA was extracted from tail clips and genotyped at Neogen (Lansing, MI) using miniMUGA arrays containing 11,000 SNP probes. DNA from parental strains and F1 hybrids was included on each array as controls. SNP markers were excluded if they showed discordant genotypes for control samples or deviated significantly from the Hardy–Weinberg equilibrium. Potential genotyping errors were further checked using the “calc errorlod” function in the R/qtl package. After filtration, 2594 informative SNPs remained for QTL analysis.

### Statistical analysis

QTL analysis was performed using the R/qtl package as previously described ^26^. Genome-wide thresholds for significant (p < 0.05) and suggestive (p < 0.63) linkage were determined by 1,000 permutations using the expectation-maximization (EM) algorithm across the genome ^27,28^. Because many traits differed significantly between male and female mice, QTL scans were performed using all F2 mice of both sexes with sex included as either an additive or an interactive covariate to maximize statistical power.

The Student t test was used to compare quantitative traits between male and female F2 mice, and linear regression was used to assess associations between two continuous variables. For traits that differed significantly by sex, the linear regression models were adjusted for sex. Statistical significance was defined as p < 0.05.

### Causal inference for atherosclerosis and body weight

Potential causal relationships between atherosclerosis and body weight were evaluated using a regression-based conditioning method ^29^. Briefly, residuals were generated from the linear regression of one trait against the other and then subjected to a genome-wide QTL scan using the same mapping algorithm. If one trait causally drives the other, conditioning on the causal mediator will eliminate or significantly reduce the LOD score of one or more QTLs for the dependent trait, rendering the residual variation independent of the driver trait.

### Candidate gene prioritization

Bioinformatic resources were used to prioritize candidate genes for the chromosome 15 (Chr 15) atherosclerosis QTL that demonstrated genetic connections with body weight. Positional candidate genes were identified based on the presence of one or more nonsynonymous coding variants or variants within upstream regulatory regions (2 kb upstream of the transcription start site) between the LP and BALB parental strains. Strain-specific variant analysis was conducted using public databases, including the Mouse Phenome Database and Ensembl. The functional impact of nonsynonymous variants on protein function was predicted using SIFT (Sorting Intolerant From Tolerant) scores.

## Results

### Sex difference in carotid atherosclerosis

Atherosclerotic lesion sizes in the left carotid arteries of female and male F2 mice were measured after 12 weeks on the Western diet. Male mice developed slightly larger lesions than female F2 mice (26,426 ± 20,057 vs. 25,201 ± 21775.8 µm^2^/section), though the difference was not statistically significant (p = 0.62) (Fig. 1). Lesion sizes varied widely among individual F2 mice, and 6% of the cohort developed no lesions (Fig. 2). Log-transformed lesion sizes followed a bimodal distribution: a bar on the far left represents the lesion-free mice, while a bell-shaped histogram on the right tracks those that developed various sizes of carotid lesions.

**Figure 1.**
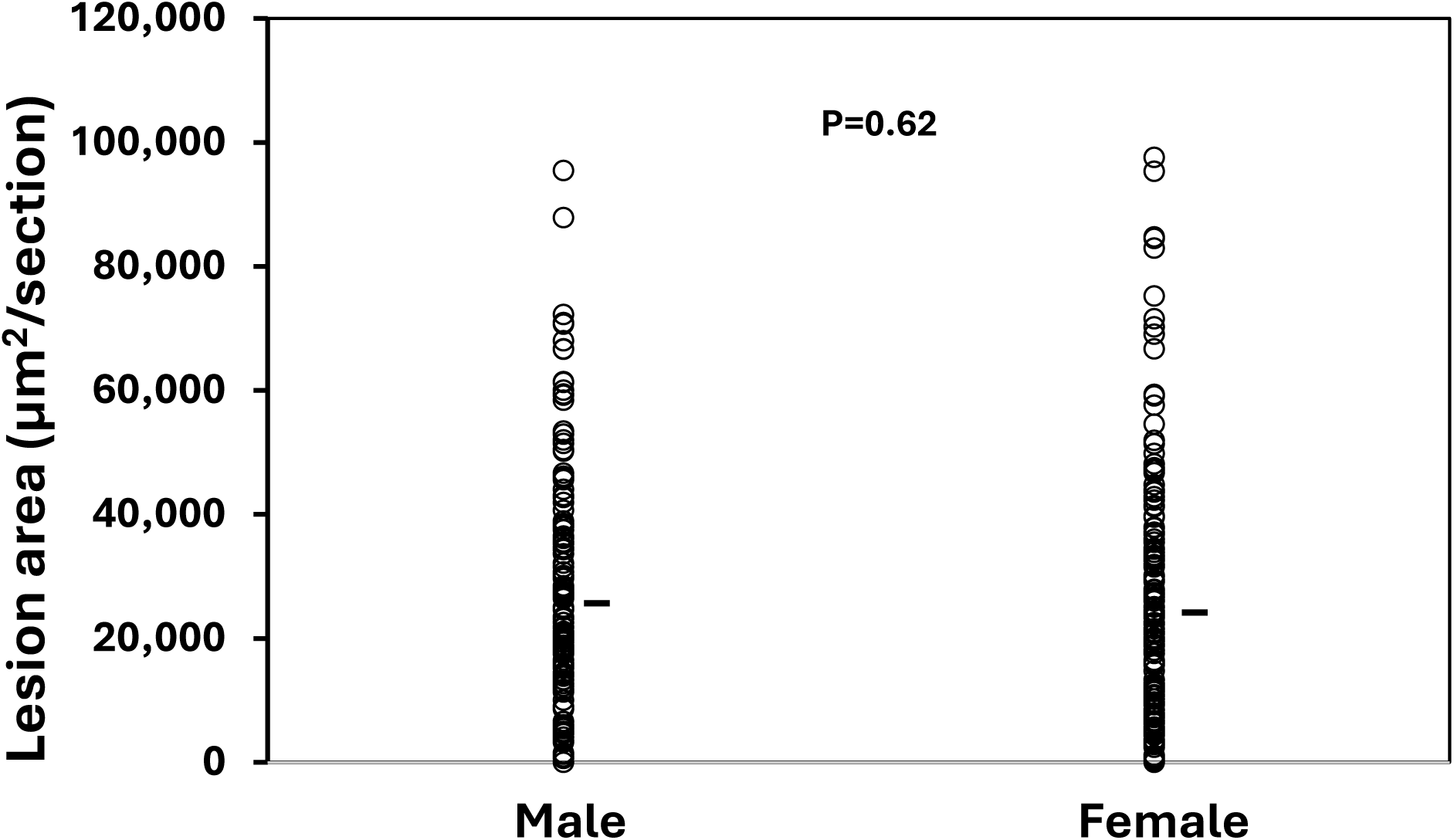
Atherosclerotic lesion sizes in the left carotid arteries of F2 mice derived from BALB/c- and LP-*Apoe*^-/-^ mice. Male and female F2 mice were fed a Western diet for 12 weeks. Each symbol represents an individual mouse. The short horizontal lines denote the mean for each group. The p value was calculated using a two-tailed Student’s T-test.

**Figure 2.**
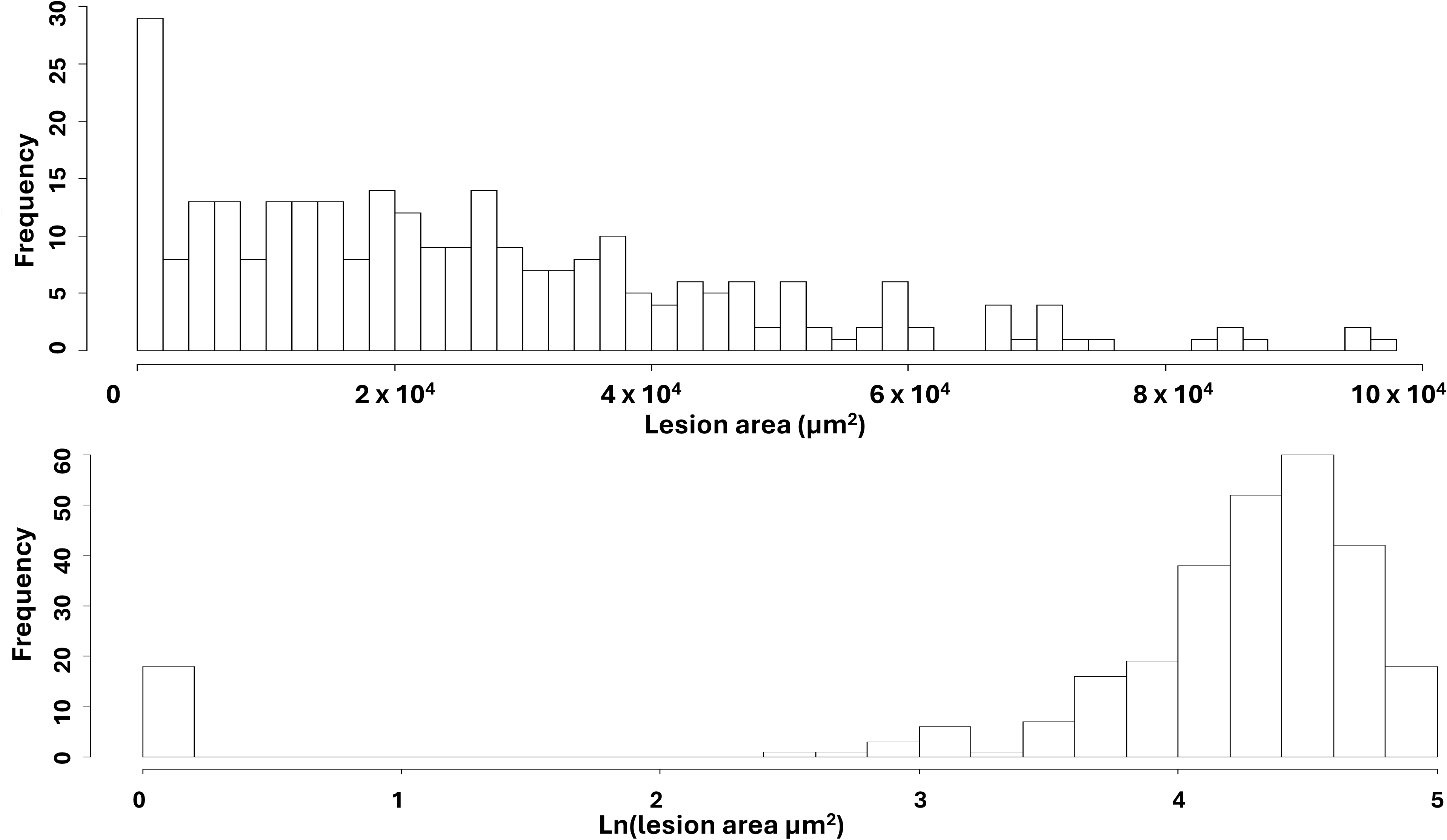
Distributions of untransformed and log-transformed carotid lesion sizes in both male and female F2 mice. Histograms showing untransformed (top) and natural log (Ln)-transformed (bottom) atherosclerotic lesion areas. Plots were generated using the plot distribution functions in R/qtl.

### QTL analysis of atherosclerosis

QTL analysis of the entire F2 cohort using sex as a covariate revealed two significant loci on chromosomes (Chr) 6 and 15, and four suggestive loci on Chr 3, 7, 13, and 19 for atherosclerotic lesion sizes (Fig. 3A). Table 1 provides comprehensive details for these loci, including locus names, LOD scores, peak markers, 95% confidence intervals (CI), modes of inheritance, high-risk alleles, and allelic effects.

**Figure 3.**
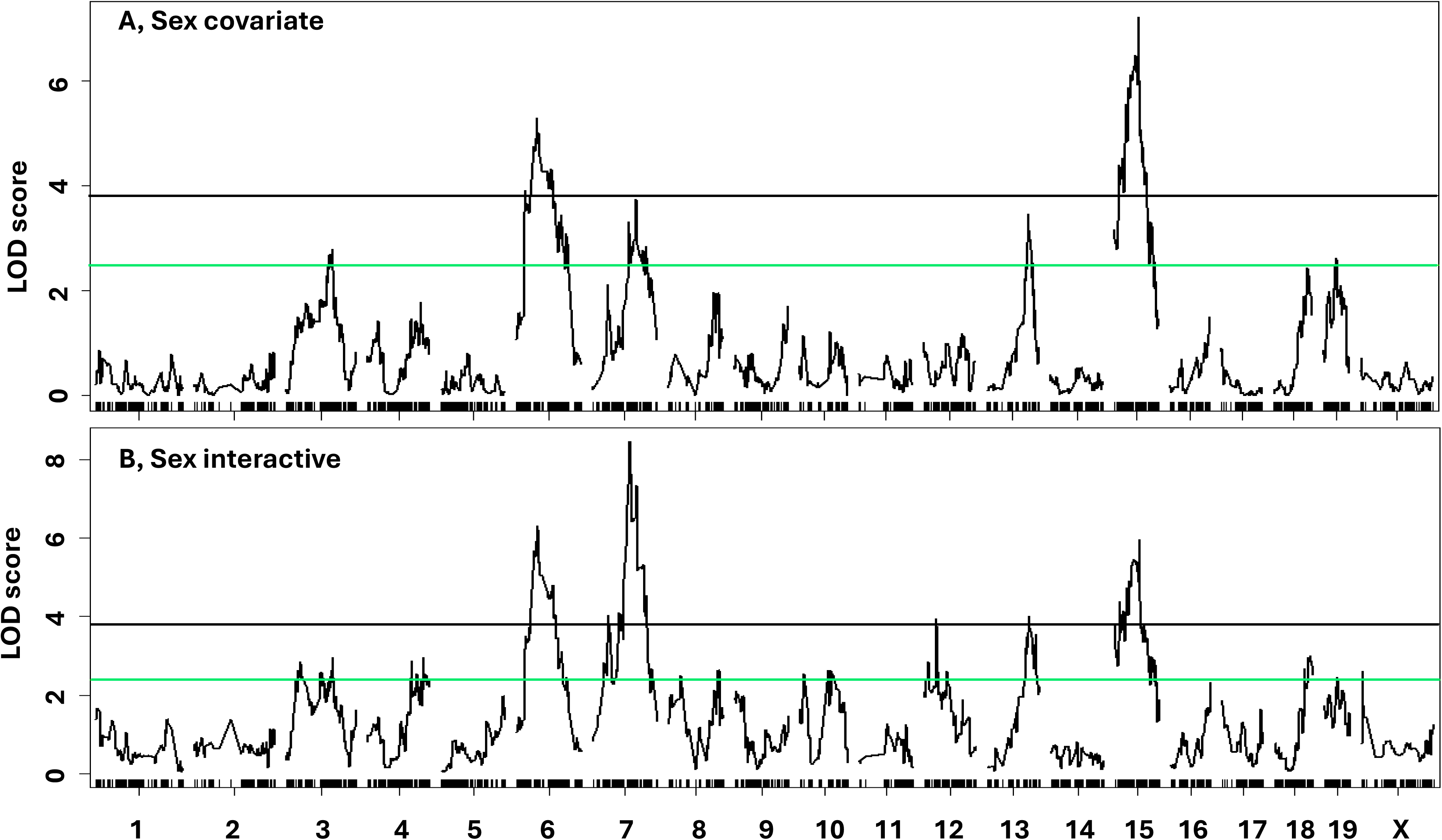
Genome-wide QTL scans for carotid atherosclerotic lesion sizes. A, Genome-wide scan across all F2 mice using sex as a covariate. B, Genome-wide scan across all F2 mice using sex as an interactive covariate. Chromosomes 1 through X are plotted along the X-axis, and the Y-axis indicates the LOD score. Horizontal green and black lines represent genome-wide thresholds for suggestive (p < 0.63) and significant (p < 0.05) linkage, respectively. Short vertical tick marks on the X-axis denote the chromosomal positions of genetic markers.

**Table 1.**
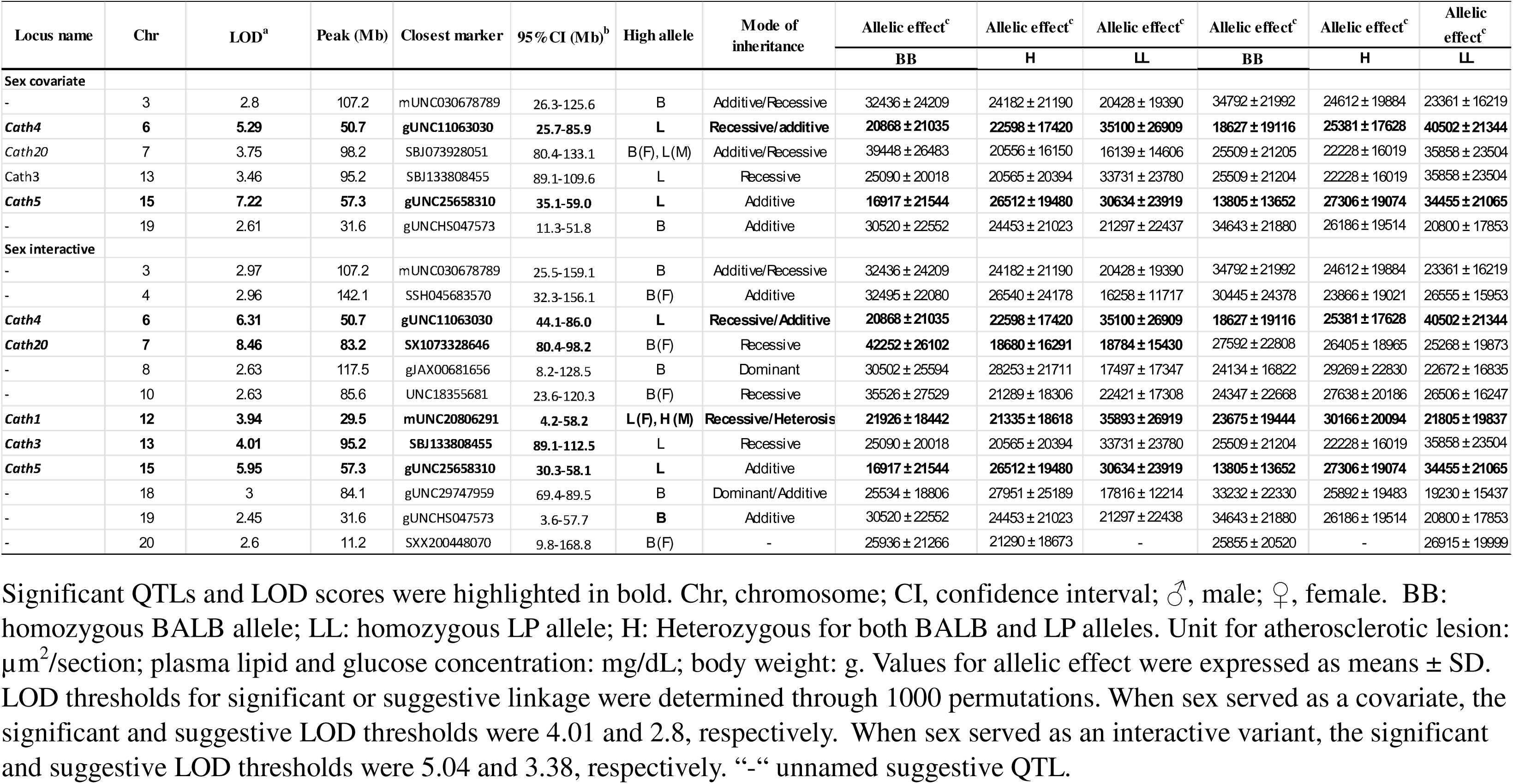
Suggestive and significant QTLs for carotid atherosclerosis mapped in F2 mice derived from LP and BALB Apoe*^-/-^* mice.

The Chr 6 QTL had a significant LOD score of 5.29 and peaked at 50.7 Mb. This QTL replicates *Cath4*, which was initially mapped in a BXH cross and subsequently replicated in a C57BL/6 x BALB *Apoe□*/□ cross ^9,10^. The Chr 15 QTL at 57.3 Mb showed a significant LOD score of 7.22, replicating *Cath5*, previously mapped in BXH and BALB x SM *Apoe□*/□ intercrosses ^30,11^. The Chr 7 QTL near 98.2 Mb and the Chr 13 QTL near 95.2 Mb had suggestive LOD scores of 3.75 and 3.46, respectively. The Chr 13 locus replicates *Cath3*, originally mapped in a C57BL/6 x BALB/c cross ^10^. The suggestive QTLs on Chr 3 and Chr 19 are novel.

We also performed a genome-wide QTL scan for carotid lesions using sex as an interactive covariate (Fig. 3B). This scan detected 5 significant QTLs on Chr 6, 7, 12, 13, and 15 and 7 suggestive QTLs on Chr 3, 4, 8, 10, 18, 19, and 20. Compared to the scan using sex as an additive covariate, the interactive sex scan revealed additional QTLs on Chr 8, 10, 12, 18, and 20. Among these, the Chr 12 QTL near 29.5 Mb was significant, with a LOD score of 3.94. This QTL replicates *Cath1*, which was initially mapped in a BXH cross and later verified in other crosses ^10,11,30^. The remaining four newly discovered QTLs are novel and showed suggestive LOD scores.

### Associations of carotid atherosclerosis with body weight and other measures

Associations of carotid lesion sizes with body weight and other measures were determined with male, female, and entire F2 cohorts (Table 2). Body weight was the only measure significantly associated with carotid lesion size in both male and female F2 mice, accounting for 5.2% of the variance in females and 4.5% in males (r^2^ values). Female F2 mice also showed significant inverse correlations with coat color intensity (r = −0.387, p = 1.6 x 10^-6^) and fasting plasma HDL cholesterol levels (r = −0.181, p = 0.03), which accounted for 15% and 3.3% of the variance in lesion sizes, respectively. In contrast, male F2 mice showed no association with these two traits or most other measures.

**Table 2.** Association of carotid lesion sizes with risk factors in F2 mice.

| Trait | R (♂+♀) | P (♂+♀) | R (♀) | P (♀) | R (♂) | P (♂) |
| --- | --- | --- | --- | --- | --- | --- |
| Body weight | **-0.140** | **0.019** | **-0.228** | **0.006** | **-0.212** | **0.013** |
| Coat color | **-0.250** | **2.60E-05** | **-0.387** | **1.56E-06** | -0.089 | 0.308 |
| LDL (chow, nonfast) | 0.067 | 0.261 | 0.138 | 0.098 | 0.014 | 0.874 |
| HDL (chow, nonfast) | 0.091 | 0.126 | 0.110 | 0.187 | 0.063 | 0.465 |
| Triglyceride (chow, nonfast) | 0.076 | 0.204 | 0.059 | 0.482 | 0.094 | 0.277 |
| Glucose (chow, nonfast) | 0.041 | 0.489 | 0.088 | 0.290 | 0.012 | 0.888 |
| LDL (Western, nonfast) | 0.035 | 0.563 | 0.035 | 0.675 | 0.099 | 0.251 |
| HDL (Western, nonfast) | 0.010 | 0.863 | -0.150 | 0.071 | 0.127 | 0.138 |
| Triglyceride (Western, nonfast) | 0.033 | 0.587 | 0.129 | 0.122 | 0.012 | 0.891 |
| Glucose ( Western, nonfast) | 0.034 | 0.572 | -0.145 | 0.082 | 0.163 | 0.058 |
| LDL (Western, fast) | 0.057 | 0.342 | 0.127 | 0.128 | 0.007 | 0.933 |
| HDL (Western, fast ) | 0.039 | 0.514 | **-0.181** | **0.030** | 0.032 | 0.712 |
| Triglyceride (Western, fast) | 0.006 | 0.924 | 0.065 | 0.448 | -0.062 | 0.479 |
| Glucose (Western, fast) | 0.100 | 0.095 | 0.105 | 0.095 | 0.099 | 0.253 |
| TBARS | **0.119** | **0.047** | 0.127 | 0.129 | 0.104 | 0.225 |
| ApoB | **0.119** | **0.046** | 0.095 | 0.259 | 0.139 | 0.104 |

Associations of atherosclerotic lesions with various risk factors were determined with generalized linear model adjusted by sex when both sexes of F2 mice were analyzed. Significant associations are highlighted in **Bold**. ♂ indicates male, ♀ female; significant correlations are highlighted in bold. Unit: body weight: gram; coat color graded into white (0), light brown (1), dark brown (2), and agouti (3); plasma lipid and glucose levels: mg/dL; MDA (malondialdehyde): $\mu$ M; small dense LDL in ApoB content: $\mu$ g/mL.

When all F2 mice were analyzed using a generalized linear model with sex as a covariate, carotid lesions were inversely associated with body weight (r = −0.14, p = 0.019) and coat color intensity (r = −0.25, p = 2.6 x 10^-5^) and positively associated with plasma malondialdehyde (r = 0.12, p = 0.047) and small dense LDL ApoB levels (r = 0.12, p = 0.046).

### Causal relationship between atherosclerosis and body weight

To evaluate the potential causal effect of body weight on carotid atherosclerosis, we performed a genome-wide QTL scan on the residuals obtained from a regression analysis of atherosclerotic lesion sizes against body weight in F2 mice. When genome-wide scan was conducted using sex as an additive covariate, the LOD score of the Chr 15 QTL decreased from 7.22 to 4.78, whereas the Chr 7 and Chr 13 QTLs were elevated from suggestive to significant linkage (Table 3, Fig. 4). The QTLs on Chr 3, 6, and 18 remained largely unaffected by body weight adjustment. Furthermore, when the residuals were analyzed using sex as an interactive covariate, the Chr 15 QTL LOD score dropped from 5.95 to 4.78, while all other QTLs remained largely unchanged (Fig. 5).

**Figure 4.**
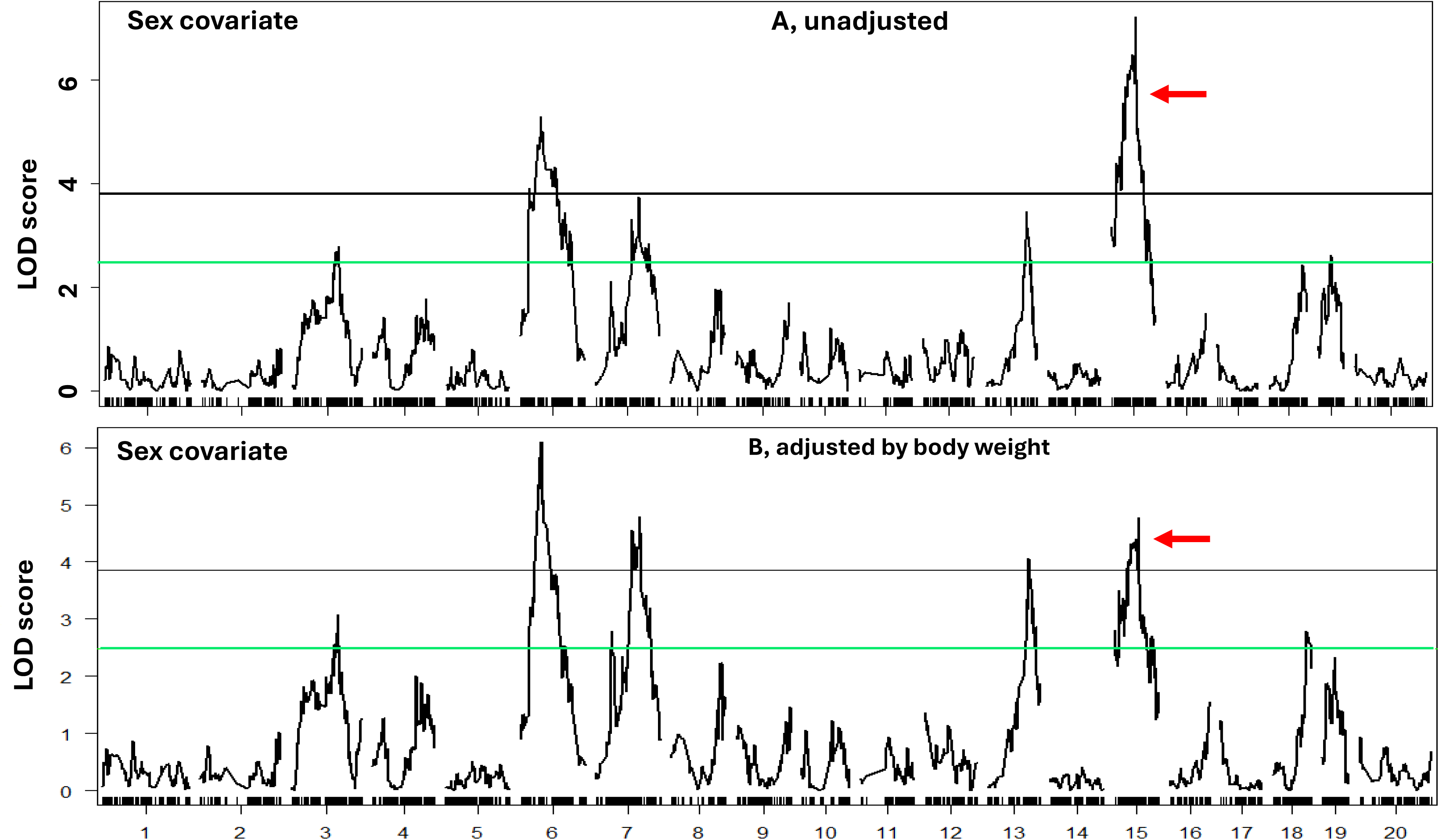
Genome-wide QTL analysis to evaluate the dependence of atherosclerosis QTLs on body weight by using sex as a covariate. **A**, Genome-wide scan for atherosclerosis without adjusting for body weight. **B**, Genome-wide scan for atherosclerosis after adjustment for body weight. Note the significant downgradation of chromosome 15 QTL (pointed by red arrow). The peak LOD scores of chromosomes 6, 7, and 13 QTLs showed moderate elevation.

**Figure 5.**
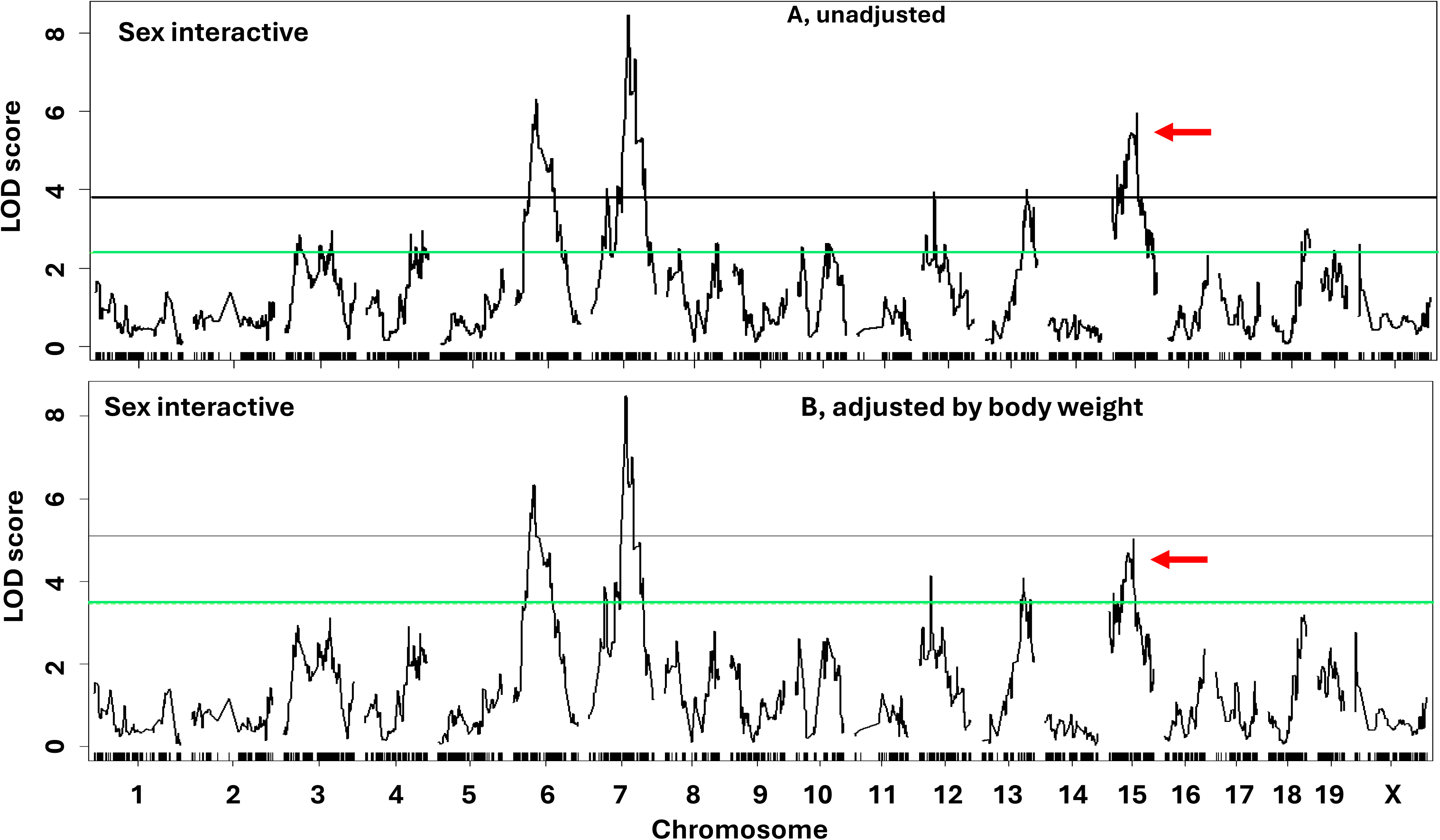
Genome-wide QTL analysis to evaluate the dependence of atherosclerosis QTLs on body weight by using sex as an interactive covariate. **A**, Genome-wide scan for atherosclerosis without adjusting for body weight. **B**, Genome-wide scan for atherosclerosis after adjusting for body weight. Note the downgradation of chromosome 15 QTL from significant to suggestive linkage (pointed by red arrow). Chromosomes 12 and 13 QTLs also downgraded from the significant to suggestive linkage, but their LOD scores showed little changes.

**Table 3.** Suggestive and significant QTLs for carotid atherosclerosis after adjusting variance in body weight.

| Locus name | Chr | LOD<sup>a</sup> | Peak (Mb) | Closest marker | 95%CI (Mb)<sup>b</sup> |
| --- | --- | --- | --- | --- | --- |
| **Nonparametric, Sex covariate** |  |  |  |  |  |
| - | 3 | 3.08 | 107.2 | mUNC030678789 | - |
| ***Cath4*** | **6** | **6.10** | **50.7** | **gUNC11063030** | **44.1-61.8** |
| ***Cath20*** | **7** | **4.79** | **98.2** | **SBJ073928051** | **79.1-114.9** |
| ***Cath3*** | **13** | **4.05** | **95.2** | **SBJ133808455** | **89.1-110.7** |
| ***Cath5*** | **15** | **4.78** | **57.3** | **gUNC25658310** | **14.2-68.0** |
| - | 18 | 2.79 | 78.7 | gUNC29665086 | - |
| **Nonparametric, Sex interactive** |  |  |  |  |  |
| ***Cath4*** | **6** | **6.33** | **50.7** | **gUNC11063030** | **44.1-86.0** |
| ***Cath20*** | **7** | **8.50** | **83.2** | **mJAX00153722** | **80.4-98.2** |
| *Cath1* | 12 | 4.13 | 29.5 | mUNC20806291 | - |
| *Cath3* | 13 | 4.08 | 95.2 | SBJ133808455 | - |
| *Cath5* | 15 | 5.03 | 57.3 | gUNC25658310 | - |

Significant QTLs and LOD scores were highlighted in bold. Chr, chromosome; CI, confidence interval. When sex served as a covariate, the significant and suggestive LOD thresholds were 3.86 and 2.89, respectively. When sex served as an interactive covariate, the significant and suggestive LOD thresholds were 5.1 and 3.45, respectively.

### Candidate genes for Chr 15 atherosclerosis QTL

Within the *Cath5* confidence interval on Chr 15, 21 protein-coding genes contain one or more non-synonymous or upstream regulatory variants between the LP and BALB strains. Among these, *Snx31, Dcstamp, Klhl38*, and *Fer1l6* contain non-synonymous SNPs that cause amino acid substitutions (Table 4). Notably, BALB mice carry two predicted deleterious mutations with low SIFT scores of 0.02: an A/G SNP (rs249728855) in *Klhl38* causing an R1450C substitution, and a G/T SNP (rs216528441) in *Fer1l6* causing a G43C substitution.

**Table 4.**
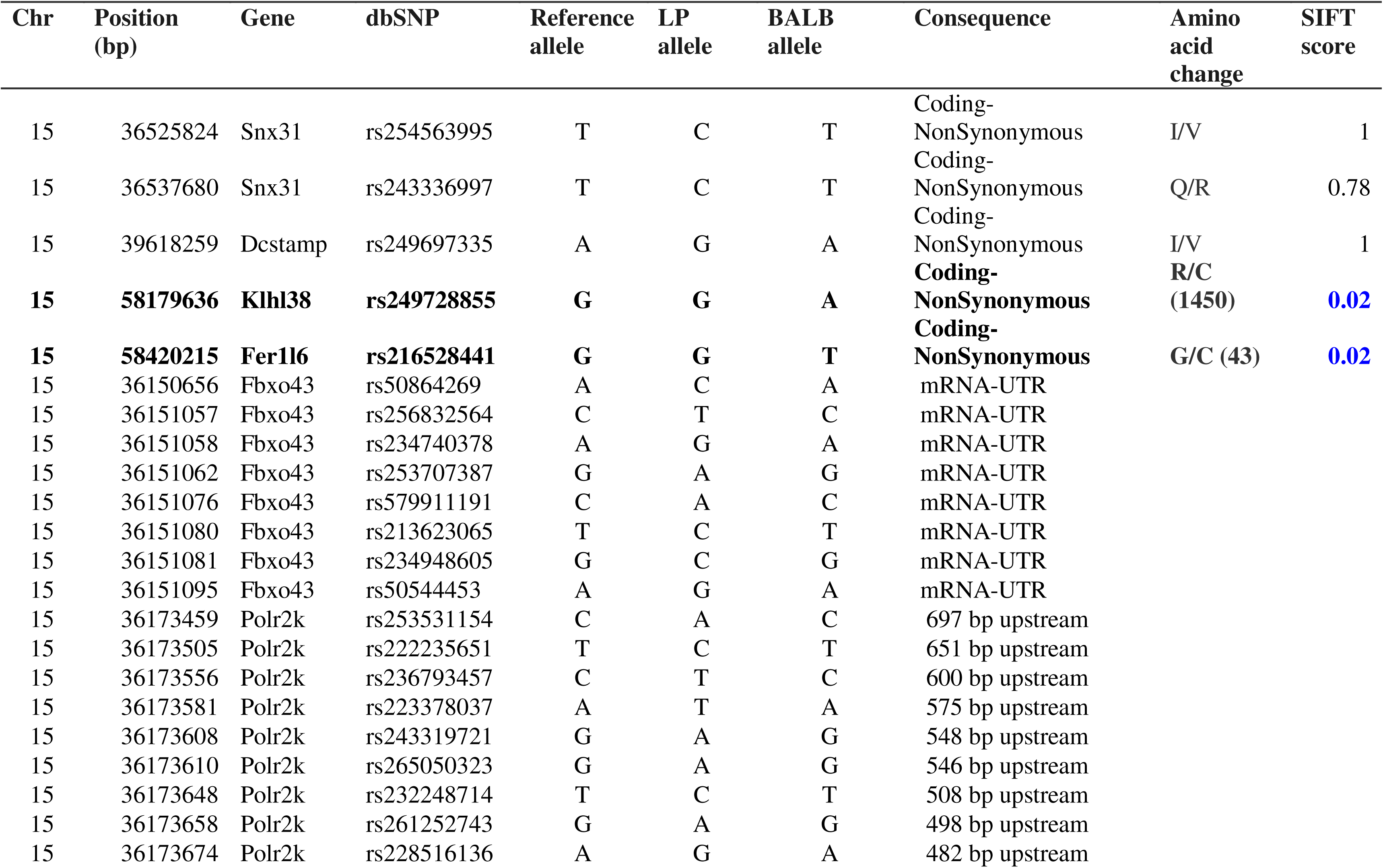
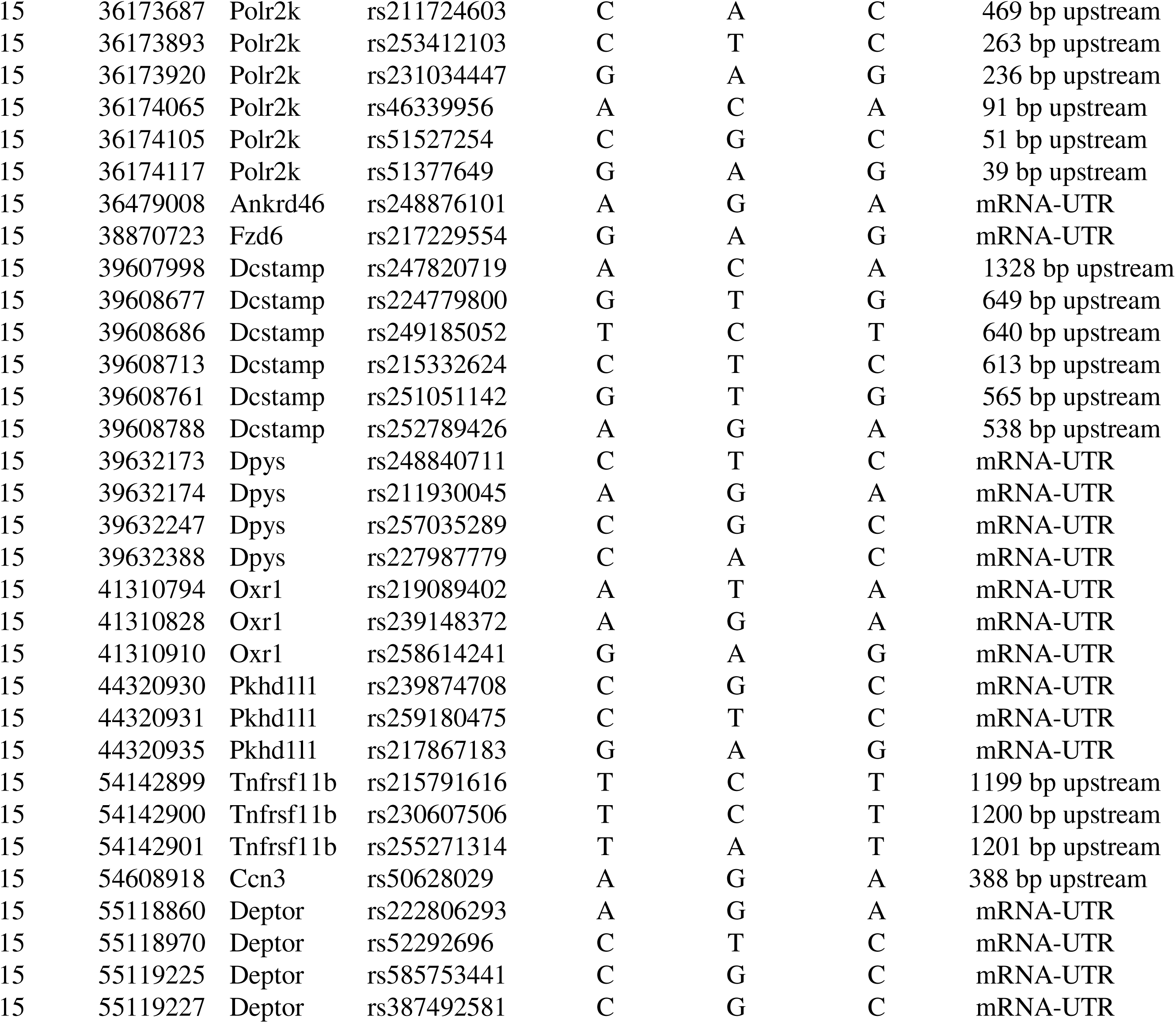
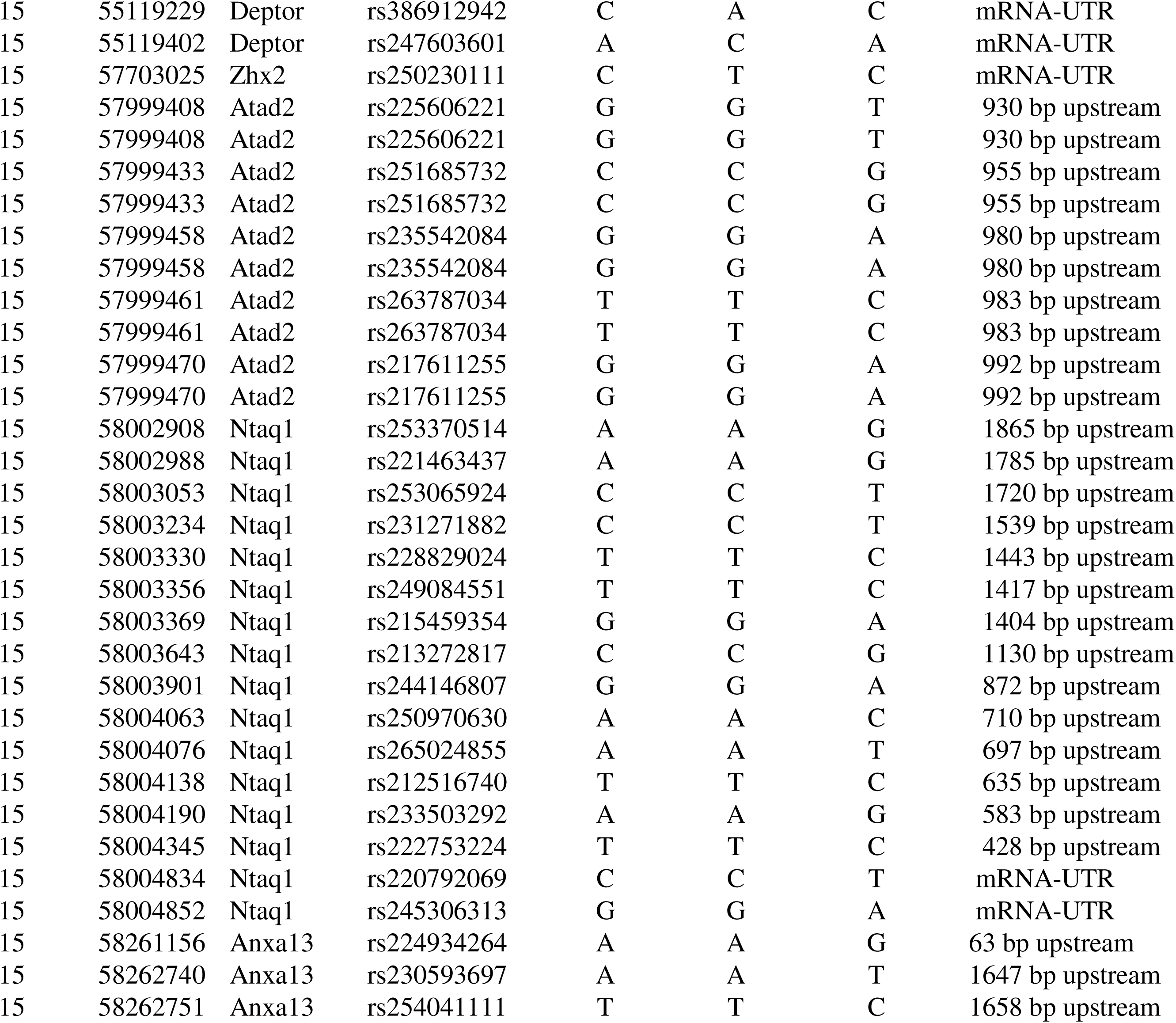
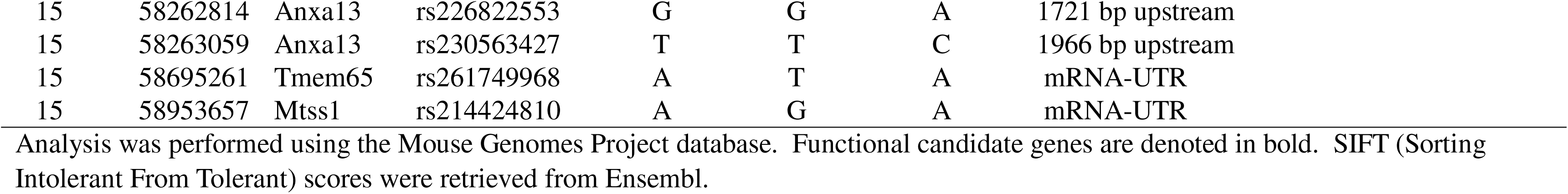
Candidate genes for *Cath5* on chromosome 15 (35–59 Mb).

| Chr | Position (bp) | Gene | dbSNP | Reference allele | LP allele | BALB allele | Consequence | Amino acid change | SIFT score |
| --- | --- | --- | --- | --- | --- | --- | --- | --- | --- |
| 15 | 36525824 | Snx31 | rs254563995 | T | C | T | Coding-NonSynonymous | I/V | 1 |
| 15 | 36537680 | Snx31 | rs243336997 | T | C | T | Coding-NonSynonymous | Q/R | 0.78 |
| 15 | 39618259 | Dcstamp | rs249697335 | A | G | A | Coding-NonSynonymous | I/V | 1 |
| **15** | **58179636** | **Klhl38** | **rs249728855** | **G** | **G** | **A** | **Coding-NonSynonymous** | **R/C (1450)** | **0.02** |
| **15** | **58420215** | **Fer1l6** | **rs216528441** | **G** | **G** | **T** | **Coding-NonSynonymous** | **G/C (43)** | **0.02** |
| 15 | 36150656 | Fbxo43 | rs50864269 | A | C | A | mRNA-UTR |  |  |
| 15 | 36151057 | Fbxo43 | rs256832564 | C | T | C | mRNA-UTR |  |  |
| 15 | 36151058 | Fbxo43 | rs234740378 | A | G | A | mRNA-UTR |  |  |
| 15 | 36151062 | Fbxo43 | rs253707387 | G | A | G | mRNA-UTR |  |  |
| 15 | 36151076 | Fbxo43 | rs579911191 | C | A | C | mRNA-UTR |  |  |
| 15 | 36151080 | Fbxo43 | rs213623065 | T | C | T | mRNA-UTR |  |  |
| 15 | 36151081 | Fbxo43 | rs234948605 | G | C | G | mRNA-UTR |  |  |
| 15 | 36151095 | Fbxo43 | rs50544453 | A | G | A | mRNA-UTR |  |  |
| 15 | 36173459 | Polr2k | rs253531154 | C | A | C | 697 bp upstream |  |  |
| 15 | 36173505 | Polr2k | rs222235651 | T | C | T | 651 bp upstream |  |  |
| 15 | 36173556 | Polr2k | rs236793457 | C | T | C | 600 bp upstream |  |  |
| 15 | 36173581 | Polr2k | rs223378037 | A | T | A | 575 bp upstream |  |  |
| 15 | 36173608 | Polr2k | rs243319721 | G | A | G | 548 bp upstream |  |  |
| 15 | 36173610 | Polr2k | rs265050323 | G | A | G | 546 bp upstream |  |  |
| 15 | 36173648 | Polr2k | rs232248714 | T | C | T | 508 bp upstream |  |  |
| 15 | 36173658 | Polr2k | rs261252743 | G | A | G | 498 bp upstream |  |  |
| 15 | 36173674 | Polr2k | rs228516136 | A | G | A | 482 bp upstream |  |  |

|  |  |  |  |  |  |  |  |
| --- | --- | --- | --- | --- | --- | --- | --- |
| 15 | 36173687 | Polr2k | rs211724603 | C | A | C | 469 bp upstream |
| 15 | 36173893 | Polr2k | rs253412103 | C | T | C | 263 bp upstream |
| 15 | 36173920 | Polr2k | rs231034447 | G | A | G | 236 bp upstream |
| 15 | 36174065 | Polr2k | rs46339956 | A | C | A | 91 bp upstream |
| 15 | 36174105 | Polr2k | rs51527254 | C | G | C | 51 bp upstream |
| 15 | 36174117 | Polr2k | rs51377649 | G | A | G | 39 bp upstream |
| 15 | 36479008 | Ankrd46 | rs248876101 | A | G | A | mRNA-UTR |
| 15 | 38870723 | Fzd6 | rs217229554 | G | A | G | mRNA-UTR |
| 15 | 39607998 | Dcstamp | rs247820719 | A | C | A | 1328 bp upstream |
| 15 | 39608677 | Dcstamp | rs224779800 | G | T | G | 649 bp upstream |
| 15 | 39608686 | Dcstamp | rs249185052 | T | C | T | 640 bp upstream |
| 15 | 39608713 | Dcstamp | rs215332624 | C | T | C | 613 bp upstream |
| 15 | 39608761 | Dcstamp | rs251051142 | G | T | G | 565 bp upstream |
| 15 | 39608788 | Dcstamp | rs252789426 | A | G | A | 538 bp upstream |
| 15 | 39632173 | Dpys | rs248840711 | C | T | C | mRNA-UTR |
| 15 | 39632174 | Dpys | rs211930045 | A | G | A | mRNA-UTR |
| 15 | 39632247 | Dpys | rs257035289 | C | G | C | mRNA-UTR |
| 15 | 39632388 | Dpys | rs227987779 | C | A | C | mRNA-UTR |
| 15 | 41310794 | Oxr1 | rs219089402 | A | T | A | mRNA-UTR |
| 15 | 41310828 | Oxr1 | rs239148372 | A | G | A | mRNA-UTR |
| 15 | 41310910 | Oxr1 | rs258614241 | G | A | G | mRNA-UTR |
| 15 | 44320930 | Pkhd1l1 | rs239874708 | C | G | C | mRNA-UTR |
| 15 | 44320931 | Pkhd1l1 | rs259180475 | C | T | C | mRNA-UTR |
| 15 | 44320935 | Pkhd1l1 | rs217867183 | G | A | G | mRNA-UTR |
| 15 | 54142899 | Tnfrsf11b | rs215791616 | T | C | T | 1199 bp upstream |
| 15 | 54142900 | Tnfrsf11b | rs230607506 | T | C | T | 1200 bp upstream |
| 15 | 54142901 | Tnfrsf11b | rs255271314 | T | A | T | 1201 bp upstream |
| 15 | 54608918 | Ccn3 | rs50628029 | A | G | A | 388 bp upstream |
| 15 | 55118860 | Deptor | rs222806293 | A | G | A | mRNA-UTR |
| 15 | 55118970 | Deptor | rs52292696 | C | T | C | mRNA-UTR |
| 15 | 55119225 | Deptor | rs585753441 | C | G | C | mRNA-UTR |
| 15 | 55119227 | Deptor | rs387492581 | C | G | C | mRNA-UTR |
| 15 | 55119229 | Deptor | rs386912942 | C | A | C | mRNA-UTR |
| 15 | 55119402 | Deptor | rs247603601 | A | C | A | mRNA-UTR |
| 15 | 57703025 | Zhx2 | rs250230111 | C | T | C | mRNA-UTR |
| 15 | 57999408 | Atad2 | rs225606221 | G | G | T | 930 bp upstream |
| 15 | 57999408 | Atad2 | rs225606221 | G | G | T | 930 bp upstream |
| 15 | 57999433 | Atad2 | rs251685732 | C | C | G | 955 bp upstream |
| 15 | 57999433 | Atad2 | rs251685732 | C | C | G | 955 bp upstream |
| 15 | 57999458 | Atad2 | rs235542084 | G | G | A | 980 bp upstream |
| 15 | 57999458 | Atad2 | rs235542084 | G | G | A | 980 bp upstream |
| 15 | 57999461 | Atad2 | rs263787034 | T | T | C | 983 bp upstream |
| 15 | 57999461 | Atad2 | rs263787034 | T | T | C | 983 bp upstream |
| 15 | 57999470 | Atad2 | rs217611255 | G | G | A | 992 bp upstream |
| 15 | 57999470 | Atad2 | rs217611255 | G | G | A | 992 bp upstream |
| 15 | 58002908 | Ntaq1 | rs253370514 | A | A | G | 1865 bp upstream |
| 15 | 58002988 | Ntaq1 | rs221463437 | A | A | G | 1785 bp upstream |
| 15 | 58003053 | Ntaq1 | rs253065924 | C | C | T | 1720 bp upstream |
| 15 | 58003234 | Ntaq1 | rs231271882 | C | C | T | 1539 bp upstream |
| 15 | 58003330 | Ntaq1 | rs228829024 | T | T | C | 1443 bp upstream |
| 15 | 58003356 | Ntaq1 | rs249084551 | T | T | C | 1417 bp upstream |
| 15 | 58003369 | Ntaq1 | rs215459354 | G | G | A | 1404 bp upstream |
| 15 | 58003643 | Ntaq1 | rs213272817 | C | C | G | 1130 bp upstream |
| 15 | 58003901 | Ntaq1 | rs244146807 | G | G | A | 872 bp upstream |
| 15 | 58004063 | Ntaq1 | rs250970630 | A | A | C | 710 bp upstream |
| 15 | 58004076 | Ntaq1 | rs265024855 | A | A | T | 697 bp upstream |
| 15 | 58004138 | Ntaq1 | rs212516740 | T | T | C | 635 bp upstream |
| 15 | 58004190 | Ntaq1 | rs233503292 | A | A | G | 583 bp upstream |
| 15 | 58004345 | Ntaq1 | rs222753224 | T | T | C | 428 bp upstream |
| 15 | 58004834 | Ntaq1 | rs220792069 | C | C | T | mRNA-UTR |
| 15 | 58004852 | Ntaq1 | rs245306313 | G | G | A | mRNA-UTR |
| 15 | 58261156 | Anxa13 | rs224934264 | A | A | G | 63 bp upstream |
| 15 | 58262740 | Anxa13 | rs230593697 | A | A | T | 1647 bp upstream |
| 15 | 58262751 | Anxa13 | rs254041111 | T | T | C | 1658 bp upstream |
| 15 | 58262814 | **Anxa13** | rs226822553 | G | G | A | 1721 bp upstream |
| 15 | 58263059 | **Anxa13** | rs230563427 | T | T | C | 1966 bp upstream |
| 15 | 58695261 | **Tmem65** | rs261749968 | A | T | A | mRNA-UTR |
| 15 | 58953657 | **Mtss1** | rs214424810 | A | G | A | mRNA-UTR |

Analysis was performed using the Mouse Genomes Project database. Functional candidate genes are denoted in bold. SIFT (Sorting Intolerant From Tolerant) scores were retrieved from Ensembl.

Seventeen genes, including *Fbxo43, Polr2k, Ankrd46, Fzd6, Dcstamp, Dprs, Oxr1, Pkhd1l1, Tnfrsf11b, Ccn3, Deptor, Zhx2, Atad2, Ntaq1, Anxa13, Tmem65,* and *Mtss1*, contain one or more SNPs in their upstream regulatory regions, which may alter their protein expression levels.

### Association with carotid atherosclerosis and adiposity in humans

The confidence interval of *Cath5* extended from 35 to 59 Mb mouse chromosome 15, corresponding to syntenic regions on human chromosomes 5 (9–10.3 Mb) and 8 (96–125 Mb). Using GWAS meta-analysis data from the Cohorts for Heart and Aging Research in Genomic Epidemiology (CHARGE) consortium ^31^, we evaluated the associations of these syntenic regions with carotid intima-media thickness (cIMT) and adiposity traits. Regional association plots revealed that the Chromosome 8 syntenic region was significantly associated with cIMT (rs10217064, p = 1.93 x 10^-8^) and waist-to-hip ratio (WHR) (rs16892421, p = 6.94 x 10^-7^) (Fig. 6A, B). Furthermore, this region demonstrated robust associations with HDL cholesterol (rs3824201, p = 6.35 x 10^-12^) and fasting glucose levels (rs3802177, p = 7.10 x 10^-25^) (Fig. 6C, D).

**Figure 6.**
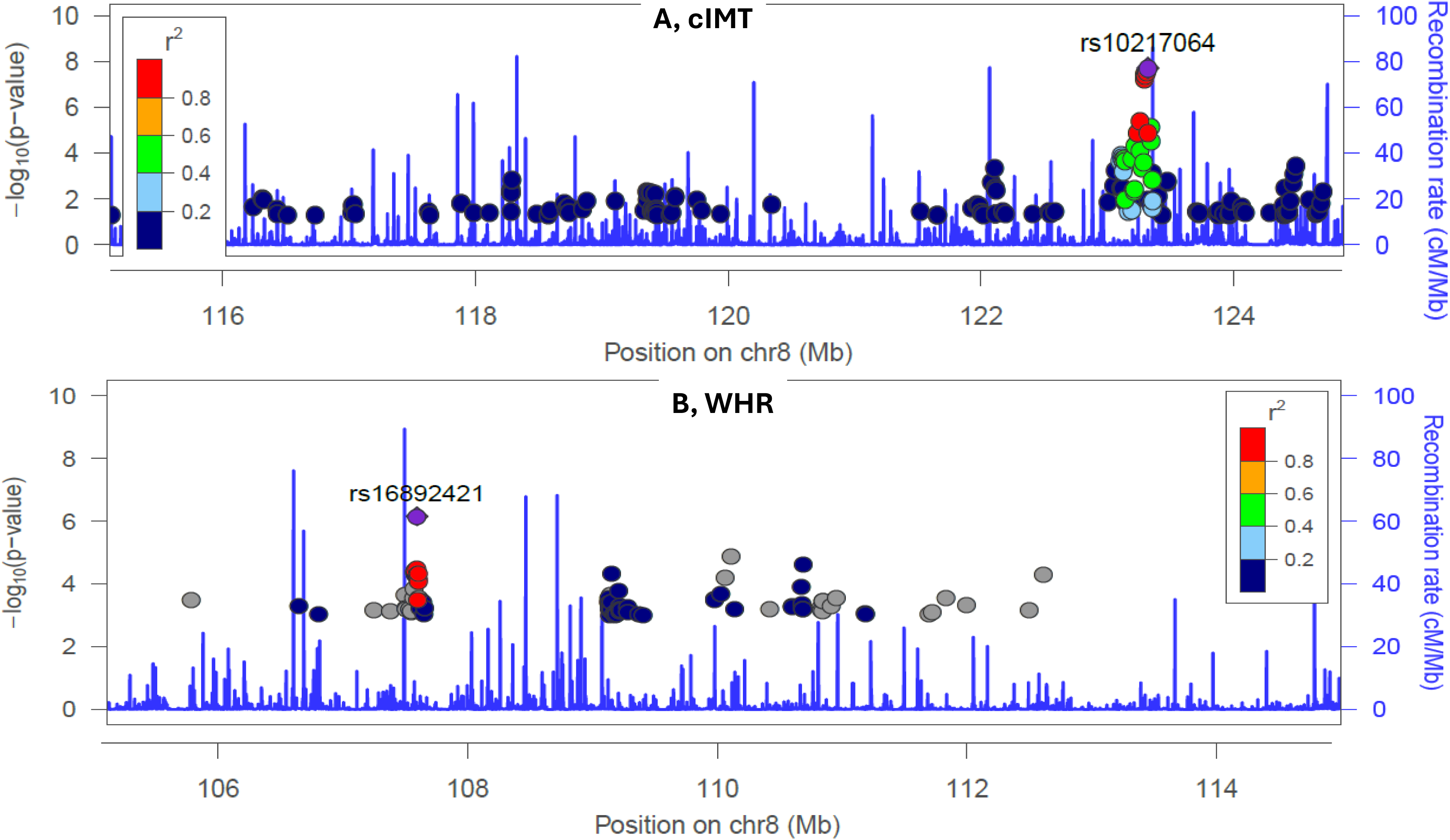
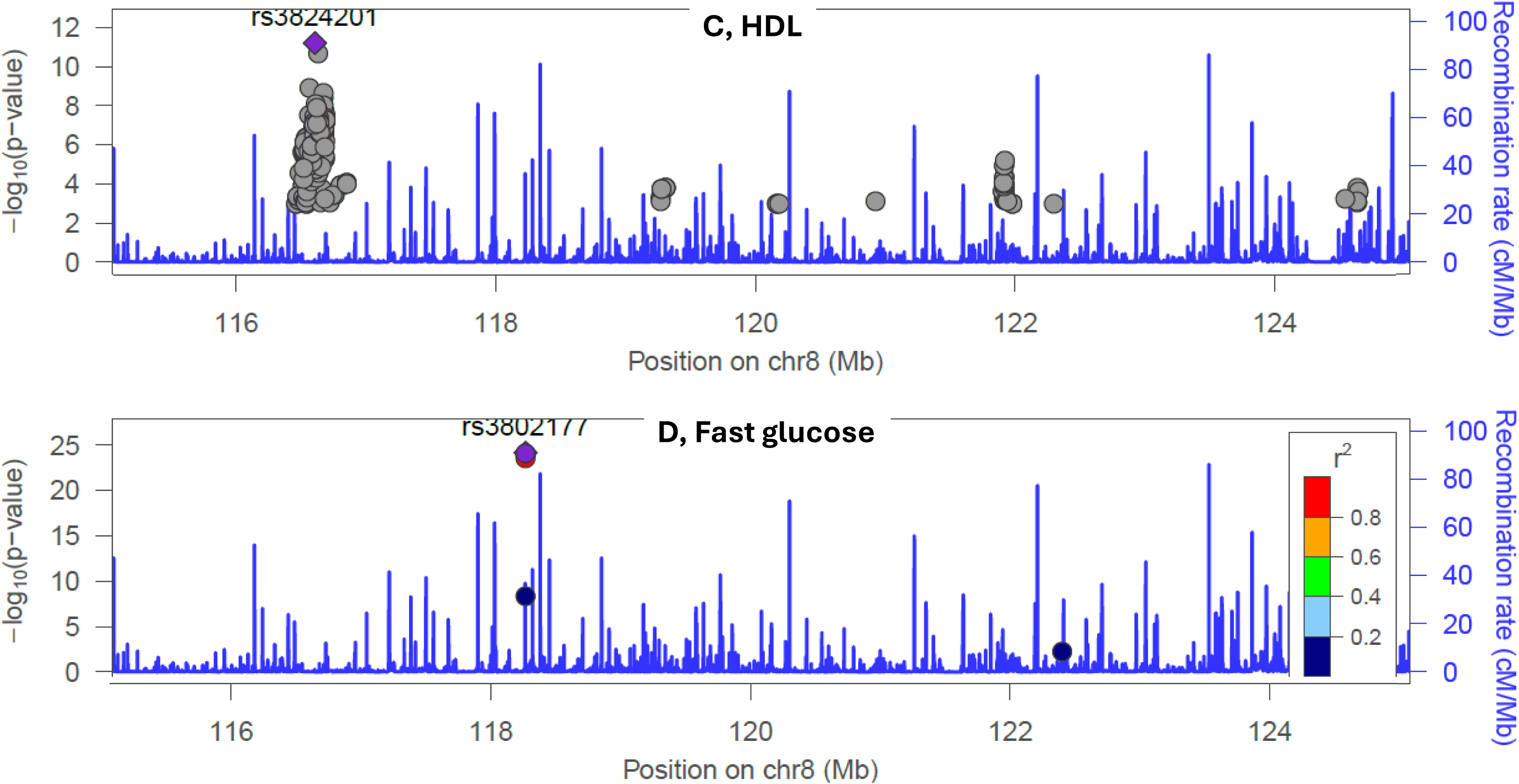
Regional association plots for human chromosome 8 (96–125 Mb), showing associations with carotid intima-media thickness (cIMT), waist-to-hip ratio (WHR), HDL cholesterol, and fasting glucose levels. GWAS meta-analysis dataset was derived from the CHARGE consortium. This genomic region corresponds to the confidence interval of *Cath5* on mouse chromosome 15.

## Discussion

In this study, we observed an inverse correlation between carotid lesion size and body weight in both male and female F2 cohorts, supporting the existence of the "obesity paradox" in carotid atherosclerosis. Two significant QTLs on chromosomes 6 and 15 were identified for carotid atherosclerosis in the additive sex model and five significant QTLs on chromosomes 6, 7, 12, 13, and 15 were mapped in the interactive sex model. Adjusting for body weight variation downgraded the chromosome 15 QTL in both additive and interactive sex models, while other QTLs were upgraded in the additive sex model and downgraded in the interactive sex model. Additionally, the human orthologous region of *Cath5* on mouse chromosome 15 associated with carotid intima-media thickness, a common measure of subclinical atherosclerosis, and adiposity.

A major finding of this study is the inverse correlation between body weight and carotid lesion size observed in both male and female F2 mice. To our knowledge, this is the first study to report an inverse correlation between carotid atherosclerosis and body weight in a mammalian model. Similar inverse relationships were observed in two of three additional F2 intercrosses tested, specifically BALB/c x C57BL/6 and BALB/c x SM/J, but not C3H x C57BL/6 (unpublished data). The aortic root is the standard site for quantifying mouse atherosclerosis. Inverse correlations between aortic lesion size and body weight or fat mass were observed in this and other cohorts reported ^15,32,33^. Body weight serves as a robust surrogate marker for total body fat in adult mice, with correlation coefficients consistently exceeding *r* > 0.90 ^15^. Thus, these collective findings demonstrate the existence of the obesity paradox in atherosclerosis, including the carotid vascular bed of mice.

In this cross, two significant QTLs for body weight mapped to chromosome 7 (*Bw1n* or *Bdwtq*) and chromosome 15 (*Bsbob5*) ^15^, overlapping the carotid atherosclerosis loci *Cath20* and *Cath5* in the confidence interval, respectively. This colocalization provided a statistical means for defining causal relationship between the two traits. Conditioning on body weight attenuated the LOD score of *Cath5* on chromosome 15 in both additive and interactive sex models. Unlike *Cath5*, the remaining suggestive and significant QTLs upgraded in the additive sex model (which controls for main effects of sex) and downgraded in the interactive sex model (which allows the QTL effect to differ between males and females). These results indicate that the chromosome 15 QTL *Cath5* acts in a body weight-dependent but sex-independent manner for carotid atherosclerosis, whereas the remaining loci operate independently of body weight through sex-influenced genetic mechanisms. These results also reveal that body weigh variations previously acted as confounding variance, masking the primary effects of these loci on carotid atherosclerosis.

It is noteworthy that the downgrade of QTLs other than *Cath5* following body weight adjustment in the sex-interactive model resulted from elevated permutation-derived significance thresholds rather than reduced absolute LOD scores. Indeed, with the exception of *Cath5* on Chromosome 15, the LOD scores of all other loci remained essentially unchanged post-adjustment. Nevertheless, because of these more stringent statistical thresholds, seven previously suggestive QTLs lost significance. Taken together, these results suggest that *Cath5* acts through shared, opposing mechanisms on body weight and plaque burden, whereas the remaining QTLs operate independently of body weight via sex-influenced mechanisms. Indeed, incorporating sex as an interactive covariate unmasked additional loci on chromosomes 4, 8, 10, 12, 18, and 20. Significantly, the chromosome 12 QTL replicates *Cath1* mapped in BXH, B6 X BALB/c, ^9,10,11,30^. The identification of additional loci across the genome underscores the power of accounting for sex-specific interactions in carotid atherosclerosis.

While male F2 mice exhibited slightly larger atherosclerotic lesions in the left carotid artery compared to females, this difference was not statistically significant. Notably, lesion burden varied widely across the cohort, with 6% of mice remaining completely resistant to lesion formation despite 12 weeks on a Western diet. This contrasts with observations at the aortic root, where all F2 mice developed atherosclerotic lesions ^15^. Furthermore, female F2 mice developed significantly larger aortic root plaques than males in both the current cohort and previously crosses ^15,32,34,35^.

Carotid lesion sizes in female mice exhibited additional inverse correlations with both coat color intensity and fasting HDL cholesterol levels under Western diet feeding, whereas no such correlations were observed in males. *Tyr,* encoding tyrosinase, is the primary determinant of coat color in mice and represents a strong candidate gene for *Cath20* on chromosome 7. In BALB/cJ mice, a point mutation renders the tyrosinase protein nonfunctional. Tyrosinase catalyzes the oxidation of phenols (including L-tyrosine, L-DOPA, and dopaquinone) and is abundantly expressed in atherosclerotic lesions, particularly by vascular smooth muscle cells. Its inverse correlation with plaque sizes suggests a protective role in carotid atherosclerosis, although the mechanisms underlying this female-specific effect remain to be elucidated.

Across the entire cohort, carotid lesion sizes showed a weak but statistical significant correlation with plasma levels of malondialdehyde (MDA), a systemic oxidative stress marker, and small dense LDL ApoB concentrations. Other traditional risk factors, including non-HDL cholesterol, triglyceride, and glucose levels showed no association with carotid lesion sizes in the F2 mice. Poor or no associations of metabolic traits with carotid lesion sizes have been observed in other mouse crosses ^9,10,11,30^.

The mouse *Cath5* region maps syntenically to human chromosomes 5 (9–10.3 Mb) and 8 (96–125 Mb). The chromosome 8 syntenic locus has consistently associated with cIMT ^36,37,38^, although the exact genes involved remain unknown. Leveraging bioinformatic tools and resources, we prioritized *Klhl38* and *Fer1l6* as promising candidate genes for *Cath5* on chromosome 15. Both genes reside beneath the linkage peak and harbor one or more nonsynonymous coding variants that have a deleterious effect on protein function. *Klhl38*, encoding Kelch-like family member 38, and *Fer1l6*, encoding Fer-1-like family member 6, represent prime functional candidates involved in ubiquitin-proteasome signaling and membrane trafficking/repair in vascular or adipose tissues. Genome-wide association meta-analyses in over 100,000 human subjects revealed associations of *Klhl38, Zh*x2, and *Fbxo32* with cIMT ^36^. *Zhx2* and *Fbxo32* are also located within the linkage peak of *Cath5* and contain upstream regulatory variants. Other candidate genes, including *Tnfrsf11b*, *Fzd6*, *Ccn3*, *Deptor*, and *Oxr1,* contain upstream regulatory variants. *Tnfrsf11b* and *Ccn3* are of particular interest due to their involvement in vascular calcification, endothelial integrity, and smooth muscle cell proliferation.

Interestingly, the same human syntenic region showed associations with waist-to-hip ratio, HDL cholesterol, and fast blood glucose. Thus, there is a possibility that genetic factors for carotid atherosclerosis may exert effects through action on these traits. Indeed, *Zhx2* was found to function as a regulator of key genes influencing lipoprotein metabolism ^39^.

In summary, we have observed the paradoxical association of body weight, a surrogate indicator of fatness, with carotid atherosclerosis in a segregating F2 mouse population. The locus *Cath5* on Chr 15 represents a major pleiotropic genetic link between carotid atherosclerosis and body weight, operating through sex-dependent and metabolic pathways. The demonstration of genetic connections between the two traits provides an explanation for their paradoxical relationship. Functional annotation prioritizes *Klhl38, Fer1l6*, and regulatory vascular genes within *Cath5*. Further validation of these prioritized candidates will help elucidate the precise cellular mechanisms underlying the obesity paradox in carotid atherosclerosis.

## Data availability

All data reported in this article are included in Supplementary materials.

